# Reconstitution of lamin assembly on nuclear pore complex-containing membranes

**DOI:** 10.1101/2025.07.28.667287

**Authors:** Ross TA Pedersen, Yinyin Zhuang, Andres V Reyes, Shou-Ling Xu, Xiaoyu Shi, Yixian Zheng

## Abstract

Intermediate filaments called lamins line the metazoan nuclear envelope and organize the nucleus and genome. Unlike actin and microtubules, purified intermediate filament proteins assemble into non-physiological structures, making it difficult to connect lamin functions to their assembly and regulation. To overcome this challenge and shed light on physiological lamin assembly mechanisms, we conducted biochemical studies of lamin-B3 endogenously present in Xenopus laevis egg extracts, which recapitulate physiological context. When we mimicked nucleoplasm conditions, which would support assembly of lamin filaments in in-tact cells, lamin-B3 assembled into higher-order structures resembling filamentous meshworks without accompanying nuclear assembly. This ectopic lamin assembly occurs on nuclear pore complex-containing membranes, but does not apparently recruit known nuclear lamina components, demonstrating that a lamin assembly process is partially separable from the rest of the nuclear lamina and nucleus. This assembly assay in the physiological context of cellular components opens the door to further dissecting nuclear lamina function in nuclear organization.

**Summary:** Mimicking nucleoplasm conditions in Xenopus laevis egg extracts triggers a lamin assembly reaction independent of nucleus assembly, giving new insights into potential lamin assembly mechanisms.

## Introduction

The nuclear lamina is a thin protein layer on the inner surface of the nuclear envelope in metazoan cells. Not only is it important for the shape and mechanics of the nucleus, but it also interacts with and organizes components of the nuclear envelope (Guo and Zheng, 2015; Guo et al., 2014; Horn, 2014) and controls gene expression and genome organization (Guelen et al., 2008; Reddy et al., 2008; Zheng et al., 2018). The major structural component of the nuclear lamina is a meshwork of intermediate filament polymers composed of protein subunits called lamins. We know surprisingly little about how the basic biochemical properties of lamins give rise to their myriad roles in the nuclear lamina. Why lamin assembles nearly exclusively under the inner nuclear envelope and whether proteins of the inner nuclear envelope are required for proper lamin assembly or organization are examples of fundamental questions that remain unanswered.

The cellular context and molecular mechanisms that control physiological lamin assembly are poorly understood because we lack good in vitro systems for studying lamin assembly. While actin and microtubules polymerize into structurally similar filaments either in pure form or in cells (Grange et al., 2017; Martins et al., 2021), intermediate filaments – including lamins – require cellular context to assembly properly. Intermediate filaments assembled from purified subunits in vitro are structurally polymorphic (Herrmann et al., 1996; Herrmann et al., 1999; Mücke et al., 2018). Lamins behave particularly poorly in vitro. Most isoforms transiently assemble into filaments, then aggregate laterally into non-physiological structures called paracrystals (Aebi et al., 1986; Ben-Harush et al., 2009; Heitlinger et al., 1991; Moir et al., 1991). The sole C. elegans lamin ortholog is the only notable exception; conditions have been established for its assembly into individual, stable filaments (de Leeuw et al., 2018; Foeger et al., 2006; Karabinos et al., 2003). Cytoplasmic intermediate filaments and lamins are more structurally uniform in cells. Vimentin filaments are made up of five protofibrils and have a diameter of ∼10 nm (Eibauer et al., 2024), while lamins are composed of a single protofilament ∼4 nm in diameter (de Leeuw et al., 2018; Turgay et al., 2017). Meanwhile, some mutations that cause aberrant lamin assembly in nuclei do not cause corresponding in vitro assembly defects for purified lamins (Wiesel et al., 2007), further underscoring the shortcomings of studying purified lamins. An in vitro system for studying lamins that recapitulates physiological context should shed new light on lamin assembly.

Xenopus laevis egg extracts are minimally diluted, biochemically active cytoplasm that lacks the constraint of an enclosed cell and nuclear membrane, making it easier to reconstitute and study cellular processes in vitro. They have been invaluable for understanding regulation of the cytoskeleton (Geisterfer et al., 2021) and the cell cycle (Murray, 1991). Chromatin added to interphase-arrested egg extracts assembles into nuclei with functional nuclear pore complexes (NPCs) and a lamin meshwork composed primarily of lamin-B3, an egg- and embryo-specific isoform that is abundant and assembly-competent in egg extracts (Zhang et al., 1996). Reconstitution using simpler substrates, including chromatin- and Ran-coated beads, has uncovered biochemical signals that orchestrate nuclear assembly (Heald et al., 1996; Zhang and Clarke, 2000). This approach has been less successful in studying nuclear lamina and lamin filament assembly because they are difficult to distinguish experimentally from nuclear assembly; it is impossible to discern whether an assembly mechanism of interest pertains to lamin assembly directly, or indirectly through affecting nuclear assembly.

As it stands, we are poorly equipped to understand basic lamin cell biology, let alone how lamins organize the nucleus and genome. It is increasingly clear that the nuclear lamina impacts gene expression through controlling genome organization (Zheng et al., 2018) and the transcriptional impact of chromatin states (Marin et al., 2025). Identifying sufficient physiological conditions and components that drive a lamin assembly reaction would help us to biochemically dissect how the entire nuclear envelope region, complete with heterochromatin, is assembled to carry out these functions.

## Results

### Cell- and nucleus-free reconstitution of a regulated lamin-B3 assembly reaction

To better understand the events that culminate in lamin assembly, we studied lamin-B3 endogenously present in Xenopus egg extracts. Lamin-B3 is subject to farnesylation in Xenopus oocytes and eggs (Firmbach-Kraft and Stick, 1993), but ≥95% of it is nevertheless non-membrane-associated and soluble (Lourim and Krohne, 1993). We asked whether the apparent assembly state of lamin-B3 changes under conditions that simulate nucleoplasm, which supports lamin assembly during the cell cycle.

We mimicked the conditions of nucleoplasm in crude X. laevis egg extracts by adding Ran-L43E – a mutant that behaves as Ran-GTP – or the same buffer without Ran-L43E as a control. In interphase cells, Ran-GTP is concentrated in the nucleoplasm where it releases nuclear import receptors from their cargos. It also contributes to nuclear assembly (Hetzer et al., 2000; Zhang and Clarke, 2000) by supporting NPC formation through releasing NPC components (nucleoporins) from nuclear import receptors (Sachweh et al., 2025; Walther et al., 2003). Nuclear import receptors also inhibit lamin-B3 assembly (Adam et al., 2008), leading us to wonder how releasing them from their cargos may affect lamin-B3 in egg extracts.

Epifluorescence imaging of samples stained by immunofluorescence revealed higher-order lamin-B3 structures that resemble filaments in Ran-L43E-supplemented crude extracts (Fig. S1A). However, crude extracts are not optimal for systematically investigating lamin assembly. Lamin-B3 no longer assembles in them if they are frozen and thawed and they contain heavy components that make them unsuitable for analysis by sucrose gradient sedimentation (see below). They also contain pigment granules that exploded during Stimulated Emission Depletion (STED) microscopy, precluding super-resolution imaging (Fig. S1B).

To overcome these challenges, we clarified crude egg extracts by centrifugation to create a partially clarified cytoplasm that contains a subset of cellular membranes (Romano et al., 2019). These “cleared” extracts retain the ability to form nuclei around added sperm chromatin even after being stored frozen (Fig. 1A). Using cleared egg extracts allowed us to examine reconstituted lamin-B3 assemblies with greater resolution. STED imaging revealed that Ran-L43E alone hardly triggered any lamin-B3 assembly in cleared egg extracts (Fig. 1B) as compared to what we qualitatively observed in crude egg extracts (Fig. S1A). Glycogen is a major component of crude egg extracts that is removed while making cleared egg extracts (Romano et al., 2019). We reasoned that it may function as a crowding agent that recapitulates the elevated colloid osmotic pressure of nucleoplasm (Lemière et al., 2022; Mitchison, 2019). Thus, we added an inert crowding agent, polyethylene glycol (PEG) 3350 to cleared egg extracts and found that under this condition, Ran-L43E stimulated considerable lamin-B3 assembly as visualized by STED microscopy (Fig. 1B). Co-staining with an alternative lamin-B3 antibody supported the conclusion that the observed structures are composed of lamin-B3 (Fig. S1C-D). While many of the lamin-B3 structures we observed resemble tangled meshworks, others look like isolated filaments (Fig. 1B). When we measured these putative lamin-B3 filaments, their lengths were lognormally distributed with a median of 0.937 µm (Fig. 1C). The length distribution of lamin filaments in nuclei is similarly right-skewed (Turgay et al., 2017) with some length variability between cell types (Sapra et al., 2020), suggesting that our reconstituted structures may model some aspects of physiological lamin filament assembly.

**Figure 1:**
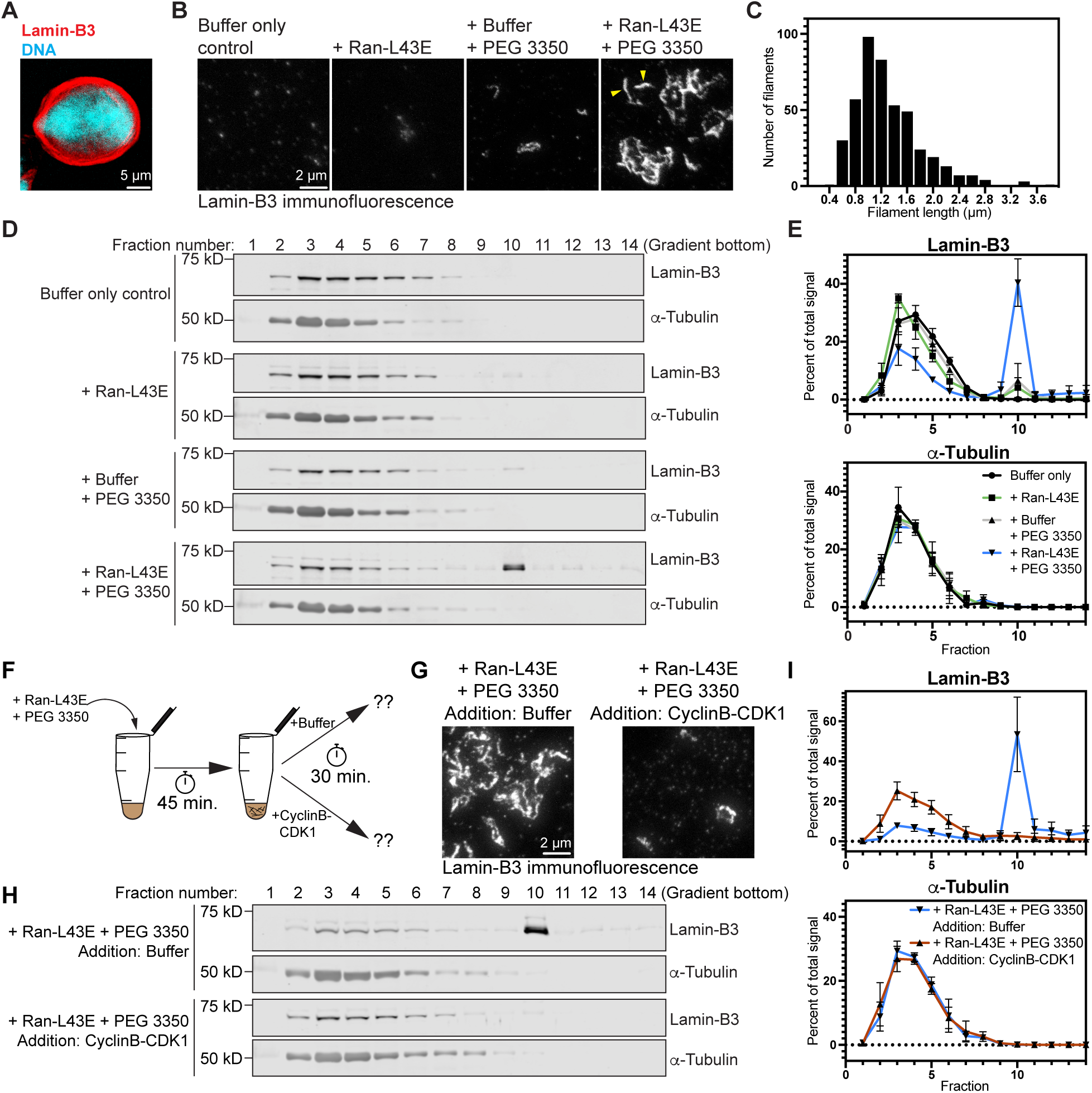
Reconstitution of regulated lamin-B3 assembly in the absence of nuclear assembly. (A) Two-color confocal slice of a nucleus assembled around demembranated sperm chromatin in “cleared” X. laevis egg extracts. Lamin-B3 in red, DNA in cyan. (B) Maximum intensity projections from z-stacks of STED microscopy lamin-B3 immunofluorescence of cleared egg extracts treated as indicated. Ran-L43E was added to ∼25 µM. Arrowheads in the Ran-L43E/PEG 3350 image highlight structures that resemble isolated filaments. (C) Lamin-B3 filament length distribution measured in extracts treated with Ran-L43E and PEG 3350 as shown in the rightmost image of panel B. Filament lengths from two biological replicate experiments were lognormally distributed, both as individual replicates and as a combined dataset, as displayed (D’Agostino and Pearson test: K^2^ = 4.35, p = 0.1136, n = 436). (D) Example immunoblots showing how lamin-B3 and α-tubulin (as a control) sediment on sucrose gradients for cleared egg extracts treated, as indicated, with the same conditions as in panel B. (E) Quantification of the fraction of total lamin-B3 and α-tubulin present in each fraction of sucrose gradients from three biological replicates of the experiment depicted in panel D. Error bars are standard deviation (SD). (F) Schematic of an experiment designed to test whether lamin-B3 assembly in our reconstitution assay is reversed by CyclinB-CDK1 function. (G) Maximum intensity-projected STED lamin-B3 immunofluorescence images of the experiment schematized in panel F. (H) Sucrose gradient sedimentation was used to assay lamin-B3 assembly state in the experiment schematized in panel F. Example immunoblots are shown. (I) Quantification (as in panel E) of sucrose gradients from three biological replicates of the experiment in panels F and H. Error bars: SD.

We next used sucrose gradient sedimentation as a semi-quantitative assay of lamin-B3 assembly in cleared egg extracts. Extract samples treated with control or lamin-B3 assembly conditions were separated down linear 5-50% sucrose gradients, then fractions from the gradient were analyzed by immunoblotting (Fig. 1D-E). Conditions that give rise to lamin-B3 assembly visible by immunofluorescence (Fig. 1B) also cause lamin-B3 to sediment into the bottom half of the gradient, primarily to the tenth fraction (Fig. 1D-E), providing independent evidence of some manner of assembly under these conditions.

To determine whether our reconstituted lamin-B3 assembly reaction is regulated by the master mitotic kinase known to disassemble both the nucleus and nuclear lamina, CyclinB-CDK1, we performed our lamin-B3 assembly reaction in egg extracts, then added either CyclinB-CDK1 or buffer as a control and incubated for an additional 30 minutes (Fig. 1F). Lamin-B3 structures incubated with buffer remained detectable by immunofluorescence after 30 minutes, while treatment with CyclinB-CDK1 for the same period resulted in lamin-B3 disassembly (Fig. 1G), as expected based on the role of CyclinB-CDK1 in triggering disassembly of lamin filaments in mitosis (Heald and Mckeon, 1990; Mall et al., 2012). Disassembly (or dissociation from larger cellular structures, see below) upon treatment with CyclinB-CDK1 was also apparent by sucrose gradient analysis, as it eliminated the population of lamin-B3 that sedimented into the tenth gradient fraction (Fig. 1H-I). Thus, our nucleus-free lamin-B3 assembly reaction is properly regulated by this major cell cycle kinase.

### Proteomics reveals nuclear pore complexes with reconstituted lamin-B3 structures

In addition to being a useful readout of lamin-B3 assembly, our sucrose gradients also partially purify assembled lamin-B3 from other extract components in our reconstituted assembly assay, which allowed us to identify proteins associated with assembled lamin-B3 using mass spectrometry. Since assembled lamin-B3 reproducibly sedimented primarily to the tenth fraction in our 5-50% sucrose gradient experiments (Fig. 1D-E, Fig. 1 H-I), we compared proteins identified in the tenth sucrose gradient fraction for extracts treated with lamin-B3 assembly conditions (1% PEG 3350, 25 µM Ran-L43E) to those identified in the same fraction for control extracts treated with buffer only, which triggers no lamin-B3 assembly as detected by immunofluorescence or sucrose gradient sedimentation (Fig. 1B, Fig. 1D-E).

We found 65 unique proteins significantly associated with lamin-B3 structures in our reconstitution assay (Fig. 2A, Table S1). As expected, the most significantly enriched protein identified was lamin-B3, and lamin-B1 was also significantly enriched (Fig. 2A, Table S1). Of the remaining unique proteins enriched in the assembled lamin-B3 fraction, 26 are nucleoporins, representing nearly every NPC component (Fig. 2B). The only nucleoporins absent in the fraction that contained assembled lamin-B3 were the cytoplasmic nucleoporin CG1 and the nucleoplasmic nucleoporin TPR, each peripheral nucleoporins that are known to associate with the NPC less stably than core nucleoporins (Rabut et al., 2004). The transmembrane nucleoporins NDC1 and GP210 were also absent, although we did identify the transmembrane nucleoporin Pom121. Another 10 of the identified proteins are related to nuclear transport or associated with NPCs (Fig. 2C). Surprisingly, no canonical nuclear lamina proteins (e.g., Lamin-B Receptor, LEM domain-containing proteins) were found associated with the reconstituted lamin-B3 structures (Table S1), despite being components of the nuclear lamina in nuclei assembled in Xenopus egg extracts (Drummond et al., 1999; Gant et al., 1999; Segura-Totten et al., 2002).

**Figure 2:**
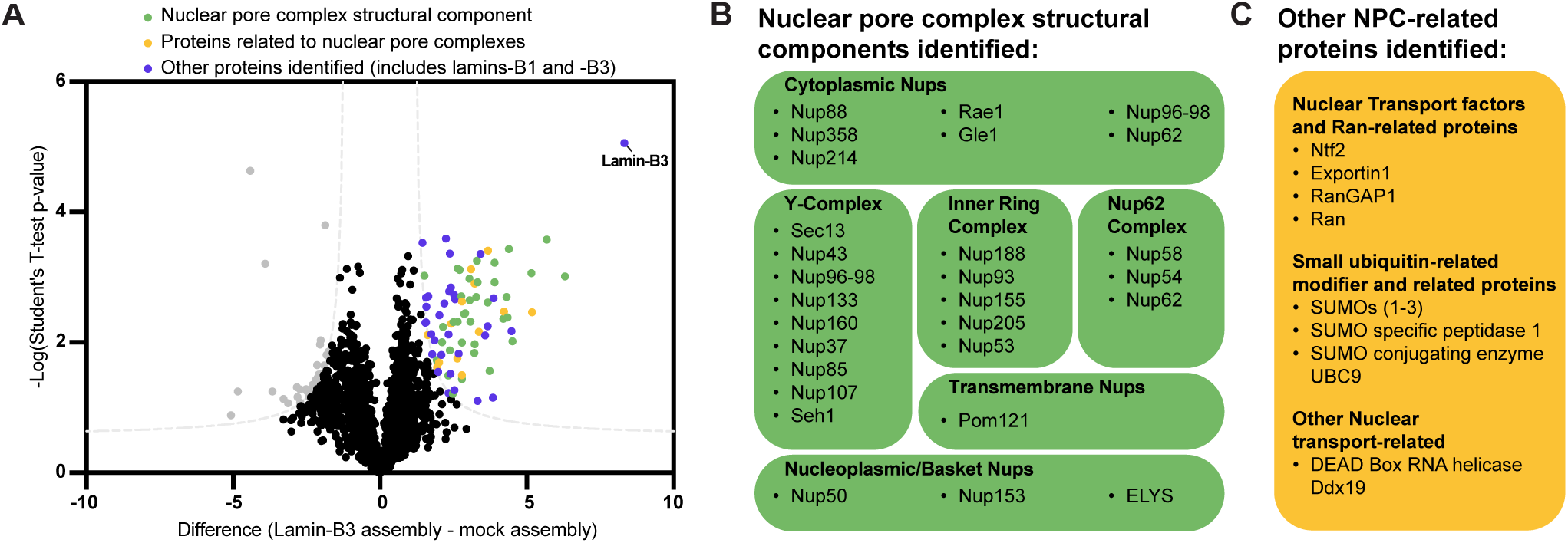
Proteomic analysis highlights the intimate relationship between lamin assembly and nuclear pore complexes. (A) Volcano plot of enrichment of proteins identified by mass spectrometry in sucrose gradient fractions that contain lamin-B3 filaments compared to proteins identified in the same fraction under mock assembly conditions as determined by two-sided T-test. Gray dashed lines are nonlinear significance thresholds (False Discovery Rate = 5%, S_0_ = 1). Gray datapoints on the left are proteins significantly enriched in mock assembly samples. Datapoints on the right are 65 unique proteins significantly enriched with assembled lamin-B3. 26 of them were nucleoporins (green). 10 of them were associated with the NPC or related to nuclear transport (yellow). Of the remaining 29 (purple), 3 were lamins, including the most significant and abundant protein identified, lamin-B3 (labeled). (B) All but 4 NPC structural proteins were found associated with lamin-B3 structures, including components from every subcomplex. (C) Examples of NPC- and nuclear transport-related proteins identified associated with lamin-B3 structures.

### Lamin-B3 assembly on nuclear pore-complex containing membranes

Lamin filaments assemble on the inner surface of the nuclear envelope and are non-randomly distributed with respect to NPCs. In interphase cells, NPCs tend to cluster within 100 nm of lamin filaments, although the two structures do not tend to colocalize (Kittisopikul et al., 2021). Our identification of so many nucleoporins in sucrose gradient fractions containing reconstituted lamin-B3 structures suggests that association of lamin-B3 with NPC-containing membranes that resemble the inner nuclear envelope may accompany this lamin assembly reaction. Our proteomic results included signatures of annulate lamellae, including nucleoporins (Fig. 2B), nuclear transport proteins (Fig. 2C; Cordes et al., 1997; Raghunayakula et al., 2015), and endoplasmic reticulum proteins (Table S1; Cordes et al., 1996). Annulate lamellae are cytoplasmic, endoplasmic reticulum-derived membrane sheets laden with NPCs. In oocytes, annulate lamellae store partially assembled NPCs to be inserted into the nuclear envelope during embryonic cell divisions (Hampoelz et al., 2016), and they have been observed in Xenopus egg extracts (Dabauvalle et al., 1991; Meier et al., 1995).

Transmission electron microscopy of membrane pellets prepared from cleared egg extracts under control and lamin-B3 assembly conditions revealed annulate lamellae as electron-dense membrane stacks (Fig. S2A). Annulate lamellae were abundant amongst the many membranes from the extracts, and they were observed at a similar density under both conditions (Fig. S2B), indicating that Ran-L43E and PEG 3350 did not cause annulate lamellae proliferation in our experiments. Two-color STED imaging of egg extract samples treated with control and lamin-B3 assembly conditions revealed that lamin-B3 assemblies decorate structures that contain NPCs (Fig. 3A) and membranes (Fig. S2C). Importantly, lamin-B3 assembly conditions caused significant enrichment of lamin-B3 signal on annulate lamellae (Fig. S2D), and visual inspection revealed that filament-like lamin-B3 structures decorate 87±6.2% of all annulate lamellae observed (Fig. 3B). Although punctate lamin-B3 immunofluorescence signal too small to reliably resolve was occasionally observed on annulate lamellae membranes under control conditions (Fig. 3A), elongated structures resolvable by STED imaging were rarely observed (Fig. 3A-B).

**Figure 3:**
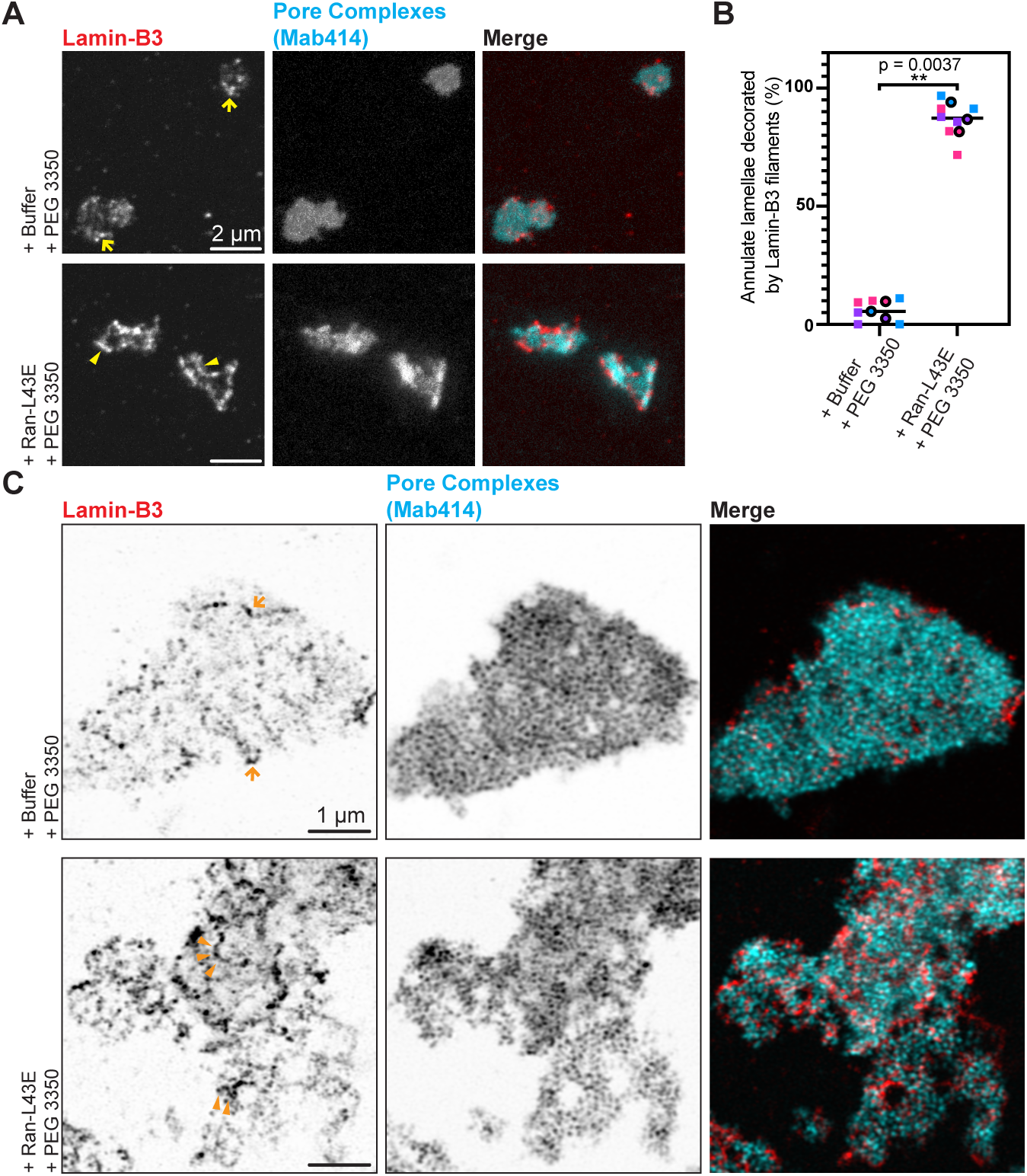
Lamin-B3 assembly occurs on annulate lamellae in the absence of nuclei. (A) Maximum intensity-projected two-color STED immunofluorescence images of lamin-B3 (red) and pore complexes (cyan, labeled by Mab414), in cleared egg extracts supplemented as indicated. Arrows in the Buffer/PEG 3350 image highlight punctate lamin-B3 signal, while arrowheads in the Ran-L43E/PEG 3350 image point to examples of filaments. (B) Quantification of the experiment from panel A: percentages of annulate lamellae decorated by lamin-B3 filaments (strong and non-diffusive staining) in the conditions indicated. Each color of square data points is from a separate biological replicate, the average for each biological replicate is displayed as an outlined circle in the corresponding color. The p value shown is from a paired two-tailed T-test comparing the biological replicate-level means (t = 16, two degrees of freedom). (C) Single-focal plane Airyscan images of expansion microscopy samples immunostained for lamin-B3 (red) and pore complexes (cyan, labeled by Mab414) in cleared egg extracts supplemented as indicated. Arrows in the Buffer/PEG 3350 image again highlight lamin-B3 signal that could be lamin-B3 oligomers or short filaments, while arrowheads in the Ran-L43E/PEG 3350 image point to examples of filaments. Scale bars correspond to 1 µm pre-expansion, with an expansion factor of 3.9.

To better resolve NPCs and lamin structures on the annulate lamellae, we applied expansion microscopy, which physically expanded samples around four times to enable imaging at a resolution of 30 nm when combined with Airyscan microscopy (Chen et al., 2015; Shi et al., 2021; Zhuang and Shi, 2023). Annulate lamellae in our samples strikingly resemble those observed by pan-expansion microscopy in HeLa cells and induced pluripotent stem cells (Morgan et al., 2025), with individual NPCs resolved. As in our STED imaging experiments (Fig. 3A), treating extracts with control conditions (buffer and PEG 3350) resulted in low-intensity, sometimes punctate lamin-B3 signal on annulate lamellae. When extracts were treated with Ran-L43E and PEG 3350, annulate lamellae were labeled with more intense lamin-B3 structures, some of them resembling serpentine filaments on the order of 1 µm in length even in single focal plane images (Fig. 3C), confirming our assembly conditions cause lamin-B3 to assemble on annulate lamellae.

Our data uncover conditions sufficient to drive lamin-B3 assembly on NPC-containing membranes that may mimic the inner surface of the nuclear envelope. These conditions include the presence of Ran-GTP and a crowding agent (perhaps simulating elevated colloid osmotic pressure or increasing the effective concentration of lamin subunits), but they intriguingly do not include chromatin, nuclear transport, and other nuclear features. Therefore, some aspects of lamin and perhaps nuclear lamina assembly, both key components of the nucleus, can be separated from the rest of nuclear assembly.

### Manipulation of the nuclear envelope milieu

To further interrogate the relationship between NPC-containing membranes and the observed lamin assembly reaction, we asked whether the outside surface of nuclei could also recruit lamin-B3 if we ectopically introduced PEG 3350 and Ran-L43E outside of assembled nuclei in the egg extracts. Indeed, ectopically introducing Ran-GTP into the cytoplasm has previously been found to give rise to nucleoplasmic structural and functional features in cytoplasm (Nachury and Weis, 1999; Sachweh et al., 2025).

We reconstituted nucleus assembly around demembranated sperm chromatin (Murray, 1991) in our cleared extracts in the presence of GST-GFP-NLS to label intact nuclei, then blocked further nucleocytoplasmic transport with 0.1 mg/mL wheat germ agglutinin (WGA) (Finlay et al., 1987). When nuclei are assembled in Xenopus egg extracts, annulate lamellae density is greatly reduced (Dabauvalle et al., 1991), so under these conditions, the predominant NPC-containing membrane present for lamin-B3 to associate with is the nuclear envelope itself. To the pre-assembled nuclei, we added an equal volume of fresh cleared extract, 1% PEG 3350, and either buffer or 25 µM Ran-L43E, then incubated the samples for a further 45 minutes. Nuclei were immediately labeled with a lamin-B3 fluorescent probe consisting of our anti-lamin-B3 antibody bound to an Alexa Fluor 594-labeled Fab secondary antibody (Weber and Bement, 2002). Lamin-B3 labeling was performed either in the absence of detergent to limit access of the lamin-B3 probe to the outside of the reconstituted nuclei, or in the presence of 0.1% triton X-100 to allow labeling on both sides of the nuclear envelopes (Fig. 4A). Assembling the nuclei in the presence of GST-GFP-NLS allowed us to definitively identify intact nuclei in samples stained without detergent, such that the lamin-B3 probe would only have access to the outer nuclear surface.

**Figure 4:**
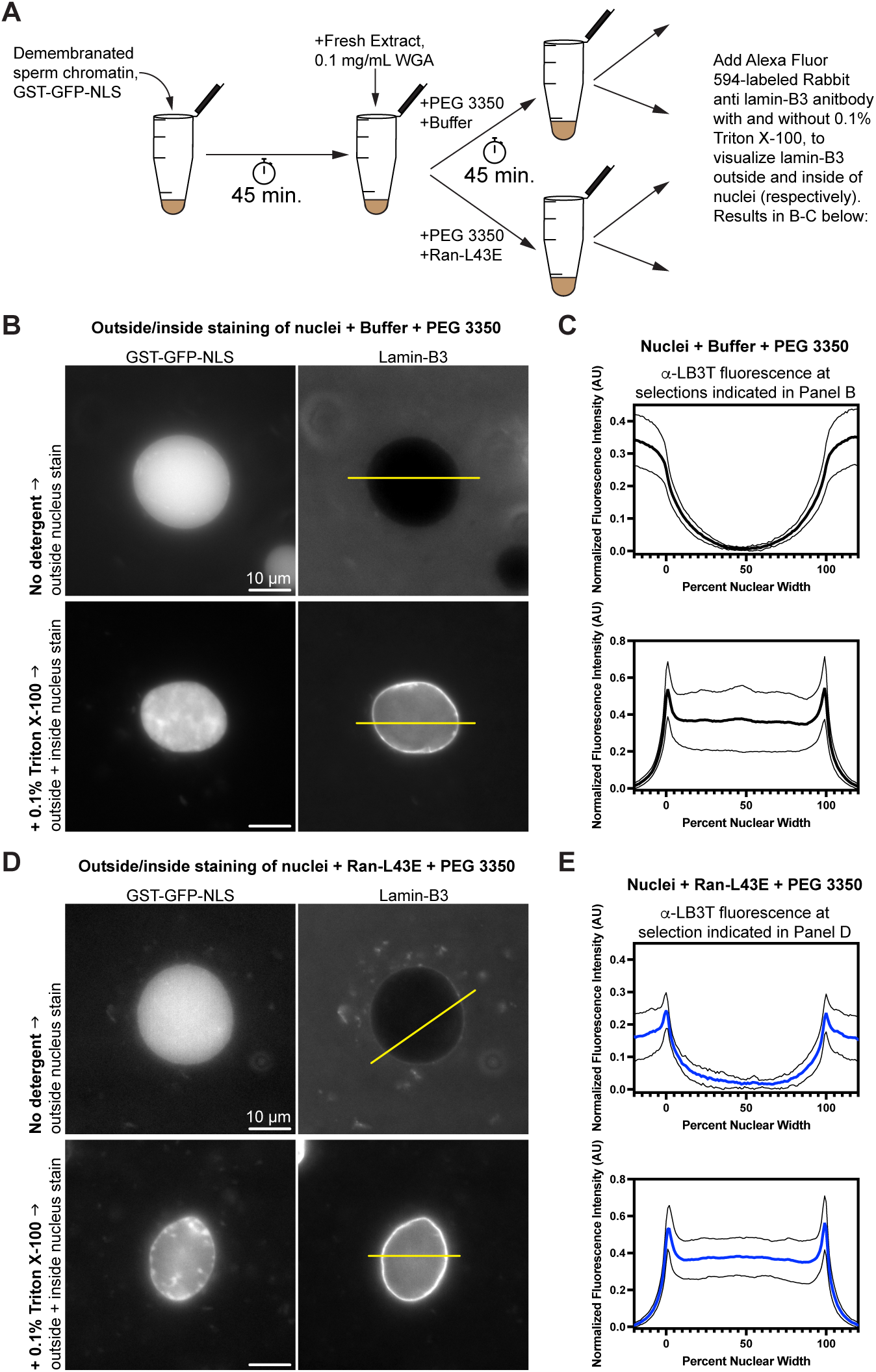
Lamin-B3 recruitment to the outside surface of the nuclear envelope. (A) Schematic of the experiment used to determine whether lamin-B3 signal comes from the outside or inside surface of the nuclear envelope. Nuclei are stained with a rabbit anti lamin-B3 antibody decorated with Alexa Fluor 594-labled goat anti rabbit fab fragments, either in the absence or presence of 0.1% triton X-100. (B) Epifluorescence images of nuclei treated with control conditions (buffer and 1% PEG 3350) and stained as illustrated in panel A. GST-GFP-NLS signal delineates nuclei. Note that nuclei stained without triton X-100 were mounted without fixation, as fixation was found to compromise nuclear integrity, but those stained in the presence of triton X-100 were fixed with formaldehyde because they otherwise become misshapen upon mounting. Fixation accounts for the non-uniform GST-GFP-NLS signal. The yellow lines are examples of selections used for intensity profile analysis like that in panel C. (C) Mean (bold lines) ± SD (fine lines) intensity profiles for lamin-B3 for the corresponding conditions in panel B. Individual intensity profiles generated from selections like those shown in panel B were normalized to their maximum value, then the minimum value was uniformly subtracted to set the baseline to zero. Mean ± SD data are from measurements on 23 (top) and 22 (bottom) nuclei collected over the course of three independent experiments. (D) Epifluorescence images of nuclei prepared as in panel B, but treated with lamin-B3 assembly conditions (25 µm Ran-L43E and 1% PEG 3350). (E) Mean (bold lines) ± SD (fine lines) intensity profiles for lamin-B3 for the corresponding conditions in panel D. Mean ± SD data are from measurements on 18 (top) and 26 (bottom) nuclei collected over the course of three independent experiments. Data were treated as those in panel C.

For assembled nuclei treated with control conditions (buffer and PEG 3350), our experiment demonstrated that lamin-B3 was only present on the inside surface of the nuclear envelope (Fig. 4B-C, top panels) because the fluorescent lamin-B3 antibody probe was excluded from the nucleoplasm and did not decorate the nuclear boundary in the absence of detergent (Fig. 4B-C). By contrast, in the presence of 0.1% triton X-100, nuclei stained brightly with the lamin-B3 probe (Fig. 4B-C, bottom panels). Therefore, lamin-B3 is only assembled on the inner surface of the nuclear envelope under this condition.

Strikingly, treating nuclei with lamin-B3 assembly conditions (Ran-L43E and PEG 3350), recruited lamin-B3 to their outside surface, as our lamin-B3 probe labeled the boundary of these nuclei even without detergent present (compare Fig. 4D-E top panels to Fig. 4B-C top panels). As expected, labeling in the presence of detergent caused a strong labeling of the nuclear rim (Fig. 4D-E, bottom panels).

This result is evidence that there may be flexibility with respect to the identity of the NPC-containing membranes where our established lamin assembly conditions can drive the assembly reaction. In circumstances where annulate lamellae are the major source of NPC-containing membranes present in cleared egg extracts, 1% PEG 3350 and 25 µM Ran-L43E drive a lamin assembly reaction on the annulate lamellae. However, under the same conditions, if nuclei and not annulate lamellae are present, lamin-B3 assembly occurs, as expected, on the inside surface of the nucleus, but lamin-B3 can also be recruited to the outside surface of the nucleus. Given that ectopically introducing Ran-GTP into the cytoplasm in other contexts gives rise to other nucleoplasmic features in the cytoplasm (Nachury and Weis, 1999; Sachweh et al., 2025), it is interesting to add lamins to the list of nuclear components that can be manipulated in this way.

## Discussion

Here, we have reported conditions sufficient to reconstitute a lamin-B3 assembly reaction that recapitulates features of physiological lamin assembly in vitro. Our results demonstrate that crowding agent and Ran-GTP drive lamin-B3, which is normally soluble in Xenopus egg extracts, to undergo an assembly reaction on NPC-containing membranes. Our experiments also reveal flexibility with respect to the identity of the NPC-containing membranes that can template this reaction, as annulate lamellae and both the inner and outer surfaces of the nuclear envelope are each able to recruit lamin-B3 under assembly conditions.

It has been speculated that purified lamins assemble filaments that self-associate laterally into paracrystals because of the absence of nuclear envelope transmembrane proteins in the “minimal” in vitro assembly system (de Leeuw et al., 2018; Herrmann and Aebi, 2016). Our results suggest that the specific nuclear envelope proteins required to preclude paracrystal formation are nucleoporins. We did not identify well-known nuclear lamina proteins associated with the lamin-B3 structures assembled in our egg extracts (Table S1), suggesting that such proteins are neither necessary for this assembly reaction to occur, nor necessarily recruited upon assembly. Biochemical interactions between lamins and nucleoporins have been repeatedly reported (Chen et al., 2013; Smythe et al., 2000) and the distribution of lamin filaments and NPCs in the nucleus has been shown to be inter-dependent (Kittisopikul et al., 2021). While further studies are needed to clarify the specific roles of nucleoporins, especially those facing the nucleoplasm, in templating or organizing lamin assembly, our findings have set the stage to further dissecting the mechanism of lamin assembly both biochemically and structurally in normal and diseased states (Buchwalter, 2023).

## Materials and Methods

### Plasmids, proteins, antibodies, and reagents

Recombinant GST-Ran-L43E was expressed from Zheng lab plasmid p213 in BL21(DE3) E. coli (New England Biolabs) and purified using standard approaches as previously reported (Dasso et al., 1994; Tsai et al., 2006; Wiese et al., 2001; Wilde and Zheng, 1999). CyclinB-CDK1 was purchased from Millipore Sigma. Lamin-B3 was visualized in blots and by immunofluorescence using rabbit and chicken polyclonal antibodies, generation of which was described previously (Ma et al., 2009). All experiments used the rabbit-generated lamin-B3 antibody except for those presented in Fig. S1C-D, which also used the chicken-generated lamin-B3 antibody. α-tubulin was visualized in blots using the DM1A mouse monoclonal antibody from Millipore Sigma. NPCs were visualized by immunofluorescence using the mouse monoclonal antibody Mab414 from Abcam, which recognizes several nucleoporins. Secondary antibodies used for immunoblots were the IR800 goat anti-rabbit antibody and the IR680 donkey anti-mouse antibodies from Licor, along with an Alexa Fluor 647 Goat anti chicken antibody from Thermo Fisher. Secondary antibodies for immunofluorescence were the Alexa Fluor 594 goat anti-rabbit antibody from Thermo Fisher (for lamin-B3 one-color immunofluorescence and two-color immunofluorescence in Fig. 3), the ATTO647N goat anti-mouse antibody from Rockland (for lamin-B3/NPC two-color immunofluorescence in Fig. 3), and the Alexa Fluor 633 goat anti-rabbit and Alexa Fluor 594 donkey anti-chicken antibodies from Thermo Fisher (Fig. S1D). DiO and Hoechst were purchased from Thermo Fisher.

### Preparation of crude *Xenopus laevis* egg extracts

Our generation of egg extracts drew on three published protocols (Chang and Ferrell, 2018; Murray, 1991; Romano et al., 2019). Mature X. laevis females were injected with 50 units of pregnant mare serum gonadotropin (BioVendor) on day 1 and 25 units on day 3. To induce ovulation, the frogs were injected with 500 units of human chorionic gonadotropin (MP Biomedicals) and placed in 2 L of 1x MMR (5 mM HEPES, 100 mM NaCl, 2 mM KCl, 2 mM CaCl_2_, 1 mM MgCl_2_, 0.1 mM EDTA, pH 7.8) on any day from day 5 to day 12. The next day, eggs were collected and rinsed four to five times with 1x MMR. Jelly coats were removed by treating for 5 minutes with 2% w/v cysteine made up in 100 mM KCl, 0.1 mM CaCl_2_, 1 mM MgCl_2_, pH adjusted to 7.8 with NaOH. Dejellied eggs were quickly rinsed four to five times with 0.2x MMR, then then activated by treating with 0.5 µg/mL calcium ionophore A23187 (Sigma-Aldrich) for 2 minutes. The eggs were then washed five times with extract buffer (“XB:” 100 mM KCl, 0.1 mM CaCl_2_, 1 mM MgCl_2_, 10 mM HEPES, 50 mM Sucrose, pH 7.7), twice with XB+ (XB with 10 µg/mL leupeptin, pepstatin, and chymostatin), and transferred to ultra-clear 13 x 51 mm ultracentrifuge tubes (Beckman). 15 minutes after adding the ionophore, the eggs were packed in a Damon/IEC Division HN-SII centrifuge by setting the speed control to “full,” spinning until the speed reached 1500 rpm, then immediately turning the instrument off and allowing the rotor to stop on its own. The packed eggs were placed on ice for 15 minutes, then crushed by centrifuging for 15 minutes at 10,000 rpm (∼12,000 xg) and 4°C in an SW 55 Ti rotor (Beckman). The layer composed of golden-colored cytoplasm was slowly withdrawn using a 1 mL syringe with an 18-gauge needle, then expelled into 1.5 mL microcentrifuge tubes on ice and supplemented with 10 µg/mL leupeptin, peptstatin, and chymostatin, 133 µg/mL cycloheximide, and energy mix (4 mM creatine phosphate, 0.4 mM ATP, 20 µg/mL creatine kinase, 0.4 mM MgCl_2_).

### Preparation of cleared egg extracts

2.2 mL of fresh crude extract prepared as described above was transferred into an ultra-clear 11 x 34 mm ultracentrifuge tube (Beckman) and centrifuged for 45 minutes at 50,000 rpm (∼166,000 xg) and 4°C in a TLS-55 rotor (Beckman). Following centrifugation, the extract is stratified into several major layers. From the top, they are: lipids, cytosol, light membranes, cytosol with some membranes mixed in, heavier membranes, and glycogen (Romano et al., 2019). Using a p1000 with a wide bore pipet tip, both cytosolic layers and the light membrane layer were collected and mixed. This “cleared extract” was snap frozen in 26 µL aliquots and stored in liquid nitrogen (storage at -80°C is also suitable for short periods).

### Reconstitution of nuclear assembly and lamin-B3 assembly in egg extracts

Reconstitution of nuclear assembly was achieved by adding 150,000-200,000 demembranated sperm chromatin masses in 1 µL (Murray, 1991) directly to 24 µL crude or cleared egg extract. Nuclear assembly was monitored over time by mixing 2 µL of the reaction with an equal volume of nucleus fix (25 mM HEPES pH 7.7, 500 mM sucrose, 8% formaldehyde, 10 µg/mL Hoechst 33350) on a glass slide, gently covering with a coverglass, and imaging on an epifluorescence microscope with a DAPI filter set. > 90% of the chromatin masses routinely assembled into nuclei within 40 minutes.

To trigger lamin-B3 assembly, extracts were supplemented with 25 µM Ran-L43E alone (crude extracts) or with 25 µM Ran-L43E and 1% w/v PEG 3350 (cleared extracts). A nuclear assembly reaction was routinely run in parallel as a positive control for extract quality and following the reasoning that the time course of lamin-B3 assembly would be at least as fast as nuclear assembly.

### Immunofluorescence sample preparation

We found that lamin-B3 structures inherently adhered to both normal glass coverslips and the poly-L-lysine coated coverslips. Lamin-B3 assembly reactions were pipetted onto clean #1.5H cover glasses, then an equal volume of 2x XB with 5% formaldehyde was added. Samples were fixed for 10 minutes, then quenched by adding glycine to a final concentration of 125 mM and incubating for 5 minutes. The liquid was aspirated and replaced with 3% w/v bovine serum albumin (BSA) made up in phosphate-buffered saline (PBS). After blocking for 30 minutes, samples were stained with the indicated primary antibodies at 3–6 µg/mL in PBS with 3% BSA for 30 minutes, then washed with five immediate exchanges of PBS with 0.1% v/v Igepal CA-630 (Sigma-Aldrich). For epifluorescence or STED imaging, samples were stained with secondary antibodies at 1.3 µg/mL in PBS with 3% BSA, washed with five immediate exchanges of PBS with 0.1% v/v Igepal CA-630, and mounted in Prolong Diamond (ThermoFisher).

### Expansion microscopy sample preparation

Samples were prepared as for immunofluorescence, but after primary antibody staining, the samples were incubated for 1 hour with goat anti rabbit antibody (Jackson Immunoresearch, Cat#111-005-144) custom conjugated with Alexa Fluor488 (Lumiprobe, Cat#21820) and goat anti mouse antibody (Jackson Immunoresearch, Cat#115-005-146) custom conjugated with Alexa Fluor 568 (Invitrogen, Cat#A20003), each at a concentration of 6 µg/mL in PBS with 0.1% triton X-100 (Zhuang and Shi, 2024). After two five-minute PBS washes, the samples were incubated in 100 mM sodium bicarbonate for five minutes, then incubated in 0.04% (w/v) glycidyl methacrylate (GMA) (Sigma, Cat#151238) in 100 mM sodium bicarbonate for three hours at room temperature. The GMA-modified sample was washed with PBS for five minutes three times, then incubated with monomer solution (8.6 g sodium acrylate, 2.5 g acrylamide, 0.15 g N,N’-methylenebisacrylamide, 11.7 g sodium chloride in 94 ml PBS) on ice for 5 minutes. Gelation solution (mixture of monomer solution, 10% (w/v) N,N,N11,N11 Tetramethylethylenediamine (TEMED) stock solution, 10% (w/v) ammonium persulfate (APS) stock solution and water at 47:1:1:1 volume ratio) was then added and samples were incubated for another 5 minutes on ice. The sample was then gelated at 37°C in a humidity chamber for 1 hour. Following gelation, the sample was gently detached from the cover slip with a forceps in heat denaturation buffer (200 mM sodium dodecyl sulfate, 200 mM NaCl, and 50 mM Tris pH 6.8), then immersed in heat denaturation buffer for 1.5 hours at 78°C. After heat denaturation, the gelated sample was fully immersed in excess DNase/RNase-free water for 30 minutes three times before being expanded in water overnight. The fully expanded gelated sample (∼3.9 times expansion) was trimmed and transferred to a glass bottom dish for imaging.

### Fluorescence microscopy and analysis

STED imaging was carried out without temperature control on a Leica TCS SP8 STED 3X imaging system with HyD detectors on a Leica DMi8 stand with either an HC PL APO 86x/1.2 NA W motCORR objective or an HC PL APO 100x/1.4 NA OIL CS2 objective. The excitation wavelength for AlexaFluor 592 was 590 nm, with detection in the range of 609-684 nm (single channel images) or 604-641 nm (dual color images). Atto 647N was excited at 653 nm and detected from 665-740 nm. For both fluorophores, a 775 nm depletion laser was used for STED imaging. The system was controlled using Leica’s LAS X software. For filament length measurements, STED images were reconstructed in 3 dimensions using Imaris (Oxford Instruments) and individual filaments were segmented and measured. For quantification of percent of annulate lamellae decorated by lamin-B3 filaments, maximum intensity projections were generated in Fiji and scored by an individual with no knowledge of which experimental condition each image came from. For quantification of annulate lamellae signal per annulate lamellae area, annulate lamellae and lamin-B3 structures were each segmented by thresholding and intensities and areas were measured using the Analyze Particles function in Fiji. Images presented in the figures are maximum intensity projections generated using Fiji.

Expansion microscopy samples were imaged without temperature control on a ZEISS LSM 980 with Airyscan 2 using a 63x water-immersion objective (Zeiss LD C-Apochromat 63x/1.15 W Corr M27). Airyscan SR and best signal mode with 0.2 AU pinhole and 1.25 AU total detection area was used for all expanded samples. The effective lateral resolution of Airyscan microscope was measured by TetraSpeck™ Microspheres (0.1 µm, fluorescent blue/green/orange/dark red) at 138 nm. After combining with expansion microscopy, the actual lateral resolution was enhanced to ∼35 nm. The system was controlled using Zeiss’s ZEN software. Images presented in the figures are single focal planes and were generated using Fiji.

Epifluorescence imaging was carried out without temperature control on a Nikon Eclipse E800 microscope with a Nikon Plan Apo 60x/1.4 NA oil immersion objective, a Hamamatsu ORCA-Flash 4.0 LT+ sCMOS camera, and an Excelitas X-Cite 120LEDmini LED illumination system. Filter sets used were Semrock’s GFP filter set (excitation filter FF01-466/40-25, emission filter FF03-525/50-25) and Texas Red filter set (excitation filter FF01-562/40-25, emission filter FF01-62440-25). The system was controlled by Metamorph. The images were visualized and quantified in Fiji. For individual intensity profiles, linear selections four pixels wide were drawn centered on a nucleus of interest such that the total length was 140% of the nuclear width. Intensity profiles were normalized for length and averaged using a previously reported Fiji plugin (Brownlee and Heald, 2019; Pedersen et al., 2020).

### Sucrose gradient sedimentation experiments

5-50% sucrose gradients (1.25 mL total) were poured as step gradients (equal volumes of 50%, 38.75%, 27.5%, 16.25%, and 5% sucrose) in XB and allowed to diffuse into continuous gradients overnight in 11 x 34 mm polycarbonate ultracentrifuge tubes (Beckman). Extract reactions were diluted 1:1 with XB and overlayed onto the gradients, then centrifuged for 4.5 hours at 50,000 rpm (∼166,000 xg) at 4°C in a TLS-55 rotor. After sedimentation, 14 equal-volume fractions were taken from the top of the gradient using a p200 with wide-bore pipet tips. Samples from each fraction were mixed with an equal volume of 2x tris urea sample buffer (125 mM Tris pH 6.8, 6 M urea, 2% sodium dodecyl sulfate, 10% 2-mercaptoethanol), boiled for 5 minutes, and resolved on 12% polyacrylamide gels. Sedimentation patterns were visualized by immunoblotting following manufacturer-recommended procedures (Bio-Rad) and imaging on a Licor Odessey CLx infrared scanner. Band intensities were measured using the associated ImageStudio software, and the plots displayed were generated in Graphpad Prism Version 10.5.0.

### Proteomics

Three biological replicate cleared egg extract samples treated with lamin-B3 assembly conditions (25 µM Ran-L43E and 1% PEG 3350) or with control conditions (buffer only) were subjected to sucrose gradient sedimentation as described above, and the tenth fraction was processed for proteomic analysis. Samples were run < 1 cm into a precast 4-12% polyacrylamide gel (Bio-Rad) and the gel was stained with Coomassie Brilliant Blue G-250 using standard procedures to reveal the boundaries between lanes. The lanes were individually excised using a clean razor blade.

In-gel trypsin digestion was performed by washing each gel after dicing it with a clean razor blade. Gel samples were repeatedly washed with 25 mM ammonium bicarbonate/50% ACN. Afterward, the samples were reduced with 10 mM DTT, alkylated with 50 mM IAM, and dried using a speed vac. Trypsin was then added and the samples were digested overnight at 37°C. The peptides were extracted by vortexing with 50% ACN/0.1% formic acid, then desalted using C18 ZipTips (Millipore).

LC-MS/MS was carried out on a Orbitrap Eclipse Tribrid Mass Spectrometer (Thermo Fisher), equipped with an Easy LC 1200 UPLC liquid chromatography system (Thermo Fisher). Peptides were first trapped using a trapping column (Acclaim PepMap 100 C18 HPLC, 75 μm particle size, 2 cm bed length), then separated using analytical column AUR3-25075C18, 25CM Aurora Series Emitter Column (25 cm x 75 µm, 1.7 µm C18) (IonOpticks). The flow rate was 300 nL/min. Peptides were eluted by a gradient from 3 to 28 % solvent B (80 % acetonitrile, 0.1 % formic acid) over 106 min and from 28 to 44% solvent B over 15 min, followed by a short wash (15 minutes) at 90 % solvent B.

Data dependent acquisition was carried out with the following parameters. Precursor scan was from mass-to-charge ratio (m/z) 375 to 1600 (resolution 120,000; AGC 200,000, maximum injection time 50ms, Normalized AGC target 50%, RF lens(%) 30) and the most intense multiply charged precursors were selected for fragmentation (resolution 15,000, AGC 5E4, maximum injection time 22ms, isolation window 1.4 m/z, normalized AGC target 100%, include charge state=2-8, cycle time 3 s). Peptides were fragmented with higher-energy collision dissociation (HCD) with normalized collision energy (NCE) 27. Dynamic exclusion was enabled for 30s.

Peptide identification and quantification of the mass spectrometry data was carried out using MaxQuant Version 2.4.10.0 (Cox and Mann, 2008). All three control and experimental samples were analyzed simultaneously using the default settings with matching between runs turned on. Label free quantification (LFQ) was enabled with the minimum ratio count set to two. The mass spectrometry data were searched against a database of 34,809 protein sequences (Proteome ID: UP000186698, available at download.xenbase.org/xenbase/Proteomes/, RRID:SCR_003280) generated from the Xenopus laevis v10.1 genome assembly (NCBI RefSeq: GCF_017654675.1). A reverted database was used to determine the false discovery rate and set it to 1%.

To analyze the resulting proteomic data, the identified protein groups and their LFQ intensities from the MaxQuant search were loaded into Perseus Version 2.0.11.0 (Tyanova et al., 2016). Data were filtered to remove decoys, potential contaminants, and protein groups that were only identified by site (i.e., posttranslational modification). LFQ intensities were log_2_ transformed, and only protein groups that had valid values for either all three experimental samples or all three control samples were kept, an operation that eliminated non-reproducible identifications. Remaining invalid values were replaced with imputed values from a normal distribution downshifted by 1.8 with a width of 0.3 made from all matrix values. A two-sided T-test with S_0_ = 1 and FDR = 0.05 was used to determine the significance and enrichment of each protein group. The volcano plot as displayed in Fig. 2A was generated in Prism.

### Transmission electron microscopy sample preparation and imaging

Electron microscopy sample preparation and image acquisition was carried out essentially as described in (Dabauvalle et al., 1991). Cleared extract samples treated with either control conditions (buffer and 1% PEG 3350) or lamin-B3 assembly conditions (25 µM Ran-L43E and PEG 3350) were incubated for 40-60 minutes at 25°C, then fixed in 112.5 mM cacodylate (pH 7.4) and 3% formaldehyde for 5 minutes at room temperature and 25 minutes at 4°C. Fixed samples were then centrifuged for 10 minutes at 3000 xg at room temperature to pellet membranes. The pellet was washed once with 0.1 M cacodylate buffer (pH 7.4) then enrobed in 2% low melt agarose made up in 0.1 M cacodylate buffer. Agarose-enrobed pellets were washed with 0.1 M cacodylate with 50 mM glycine to quench any residual formaldehyde, then rinsed with 0.1 M cacodylate buffer and stained with 2% osmium tetroxide in 0.1 M cacodylate buffer. Osmium tetroxide-stained samples were rinsed thoroughly with water, then stained for 15 minutes with 1% uranyl acetate. Samples were again rinsed thoroughly with water, then dehydrated through a concentration series into anhydrous ethanol and embedded in araldite.

Electron microscopy images were acquired on a Hitachi HT7800 transmission electron microscope operated at 80 KeV at 2000x magnification using an AMT Nanosprint 12 camera.

## Online supplemental material

Fig. S1 shows preliminary reconstitution experiments in crude Xenopus egg extracts and experiments validating our rabbit anti-lamin-B3 antibody. Fig. S2 shows additional data showing that lamin-B3 assembly occurs on annulate lamellae membranes. Table S1 reports all proteins significantly enriched in experimental and control samples for the proteomic experiment reported in Fig. 2.

## Acknowledgements

We thank Mahmud Siddiqi for providing training and advice on fluorescence microscopy and for maintaining the equipment, Ru-Ching Hsia for assistance with electron microscopy, and Lynne Hugendubler for animal care. We thank Sara Debic for critiquing the manuscript and colleagues at the Carnegie Institution for Science Department of Embryology for feedback, encouragement, and moral support. This work was supported by the following National Institutes of Health (NIH) and National Science Foundation (NSF) grants: NIH F32GM142145 (to R.T.A.P.), NIH R01GM106023 (to Y. Zheng. and R.D. Goldman), NIH R01GM110151 (to Y. Zheng), NIH DP2GM150017 (to X.S.), and NSF CAREER Award 2341058 (to X.S.). The Carnegie Mass Spectrometry Facility is supported by NIH grant S10OD030441 (to S.-L.X.). The authors declare no competing financial interests.

---

**Figure S1:**
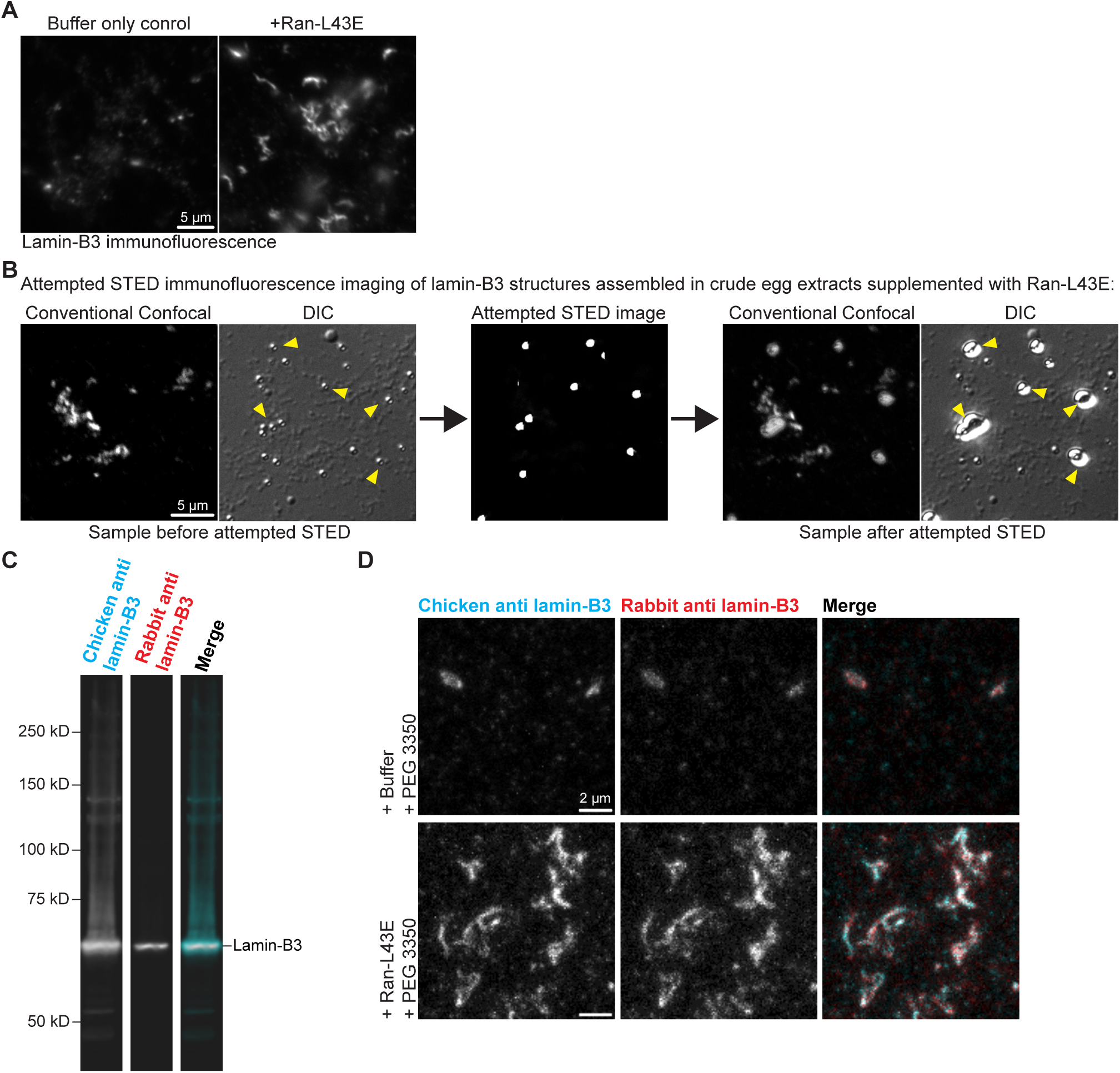
Drawbacks of reconstituting lamin-B3 assembly in crude *Xenopus* egg extracts and independent verification of anti-lamin-B3 antibody specificity. (A) Epifluorescence images of crude Xenopus egg extracts treated with a buffer control or with ∼25 µM Ran-L43E and stained by lamin-B3 immunofluorescence. Ran-L43E, but not crowding agent, is needed for lamin-B3 assembly in crude extracts, possibly because abundant glycogen present in crude extracts serves a similar role. (B) Illustration of one reason why crude extracts are not suitable for systematic studies of lamin-B3 assembly. Lamin-B3 assembly was triggered by adding Ran-L43E to crude extracts, as in panel A. Left: conventional confocal and DIC image of lamin-B3 structures before attempted STED imaging. DIC imaging reveals many small shiny granules present in the sample (arrowheads). Center: Attempted STED fluorescence image. The granules are excited by the depletion laser and saturate the detectors, masking lower intensity signals. Right: conventional confocal and DIC image of lamin-B3 structures after attempted STED imaging. The small granules explode, physically distorting the sample (arrowheads). (C) Two-color immunoblot for lamin-B3 in cleared egg extracts using two independently generated antibodies, one raised in chicken, one in rabbit. Both recognize the same band at lamin-B3’s predicted molecular weight (∼67 kD). (D) Maximum intensity projections from conventional confocal z-series images of lamin-B3 immunofluorescence for cleared egg extracts treated as indicated. The same chicken IgY and rabbit IgG antibodies from panel C were used for two-color labeling.

**Figure S2:**
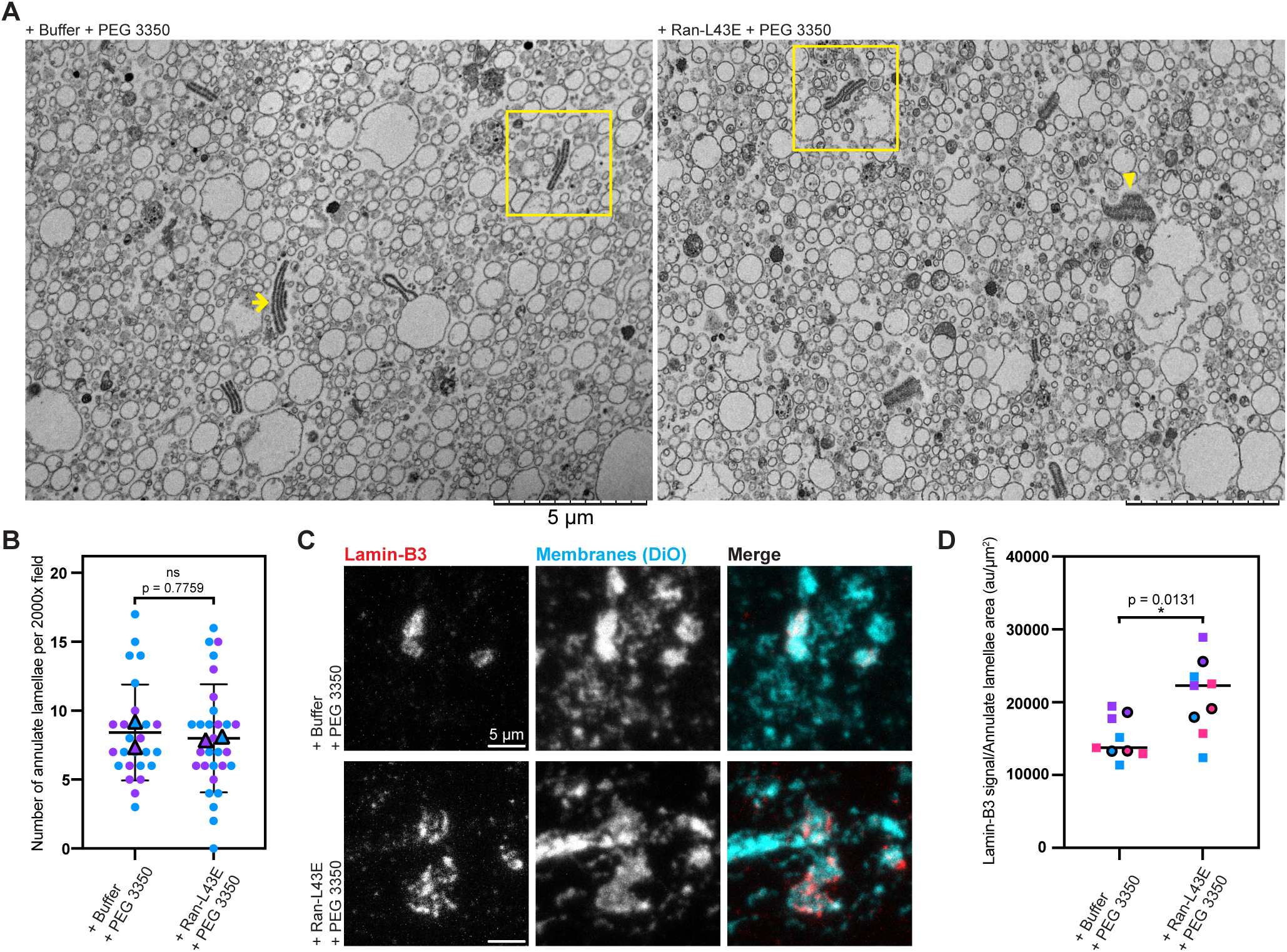
Conditions that promote lamin-B3 assembly do not alter membranes or cause annulate lamellae proliferation in cleared *Xenopus* egg extracts. (A) Transmission electron micrographs of membrane pellets prepared from cleared Xenopus egg extracts treated as indicated, stained with osmium tetroxide, resin-embedded, ultrathin-sectioned, and counterstained with uranyl acetate. One example annulate lamellae membrane stack is indicated in a yellow box in each image. Note that some annulate lamellae are captured in the section in profile such that the membrane stack is clearly visible (yellow arrow), while others are captured en face (yellow arrowhead) such that the dense pore complexes within are apparent. (B) Quantification of the density of annulate lamellae observed in images like those shown in panel A. Each color of circle data points is from a separate biological replicate, the average for each biological replicate is displayed as an outlined triangle in the corresponding color. The p value shown is from a paired two-tailed T-test comparing the biological replicate-level means (t = 0.3674, one degree of freedom). (C) Maximum intensity-projected two-color immunofluorescence images of lamin-B3 (red, imaged by STED) and membranes (cyan, labeled by DiO, imaged by conventional confocal imaging), in cleared egg extracts supplemented as indicated. (D) A second quantification of the experiment from Fig. 3A: total lamin-B3 signal per annulate lamellae area, measured in two separate images from three biological replicates. The p value shown is from a paired two-tailed T-test comparing the biological replicate-level means (t = 8.654, two degrees of freedom).

## Notes

### Competing Interest Statement

The authors have declared no competing interest.

### Summary of Updates

The revised manuscript contains many textual edits in addition to some figure reorganization and the addition of figure panels recapitulating a key result from the study with an independently-generated antibody.

## References

Adam, S. A., Sengupta, K. and Goldman, R. D. (2008). Regulation of nuclear lamin polymerization by importin α. J. Biol. Chem. 283, 8462–8468.

Aebi, U., Cohn, J., Buhle, L. and Gerace, L. (1986). The nuclear lamina is a meshwork of intermediate-type filaments. Nature 323, 560–564.

Ben-Harush, K., Wiesel, N., Frenkiel-Krispin, D., Moeller, D., Soreq, E., Aebi, U., Herrmann, H., Gruenbaum, Y. and Medalia, O. (2009). The Supramolecular Organization of the C. elegans Nuclear Lamin Filament. J. Mol. Biol. 386, 1392–1402.

Brownlee, C. and Heald, R. (2019). Importin α Partitioning to the Plasma Membrane Regulates Intracellular Scaling. Cell 176, 805–815.e8.

Buchwalter, A. (2023). Intermediate, but not average: The unusual lives of the nuclear lamin proteins. Curr. Opin. Cell Biol. 84,.

Chang, J. B. and Ferrell, J. E. (2018). Robustly cycling Xenopus laevis cell-free extracts in Teflon chambers. Cold Spring Harb. Protoc. 2018, 593–600.

Chen, H., Chen, X. and Zheng, Y. (2013). The nuclear lamina regulates germline stem cell niche organization via modulation of EGFR signaling. Cell Stem Cell 13, 73–86.

Chen, F., Tillberg, P. W. and Boyden, E. S. (2015). Expansion microscopy. Science 347, 543–548.

Cordes, V. C., Reidenbach, S. and Franke, W. W. (1996). Cytoplasmic annulate lamellae in cultured cells: Composition, distribution, and mitotic behavior. Cell Tissue Res. 284, 177–191.

Cordes, V. C., Reidenbach, S., Rackwitz, H. R. and Franke, W. W. (1997). Identification of protein p270/Tpr as a constitutive component of the nuclear pore complex-attached intranuclear filaments. J. Cell Biol. 136, 515–529.

Cox, J. and Mann, M. (2008). MaxQuant enables high peptide identification rates, individualized p.p.b.-range mass accuracies and proteome-wide protein quantification. Nat. Biotechnol. 26, 1367–1372.

Dabauvalle, M.-C., Loos, K., Merkert, H. and Scheer, U. (1991). Spontaneous Assembly of Pore Complex-containing Membranes (“Annulate Lamellae”) in Xenopus Egg Extract in the Absence of Chromatin. J. Cell Biol. 112, 1073–1082.

Dasso, M., Seki, T., Azuma, Y., Ohba, T. and Nishimoto, T. (1994). A mutant form of the Ran/TC4 protein disrupts nuclear function in Xenopus laevis egg extracts by inhibiting the RCC1 protein, a regulator of chromosome condensation. EMBO J. 13, 5732–5744.

de Leeuw, R., Gruenbaum, Y. and Medalia, O. (2018). Nuclear Lamins: Thin Filaments with Major Functions. Trends Cell Biol. 28, 34–45.

Drummond, S., Ferrigno, P., Lyon, C., Murphy, J., Goldberg, M., Allen, T., Smythe, C. and Hutchison, C. J. (1999). Temporal differences in the appearance of NEP-B78 and an LBR-like protein during Xenopus nuclear envelope reassembly reflect the ordered recruitment of functionally discrete vesicle types. J. Cell Biol. 144, 225–240.

Eibauer, M., Weber, M. S., Kronenberg-Tenga, R., Beales, C. T., Boujemaa-Paterski, R., Turgay, Y., Sivagurunathan, S., Kraxner, J., Köster, S., Goldman, R. D., et al. (2024). Vimentin filaments integrate low-complexity domains in a complex helical structure. Nat. Struct. Mol. Biol. 31, 939–949.

Finlay, D. R., Newmeyer, D. D., Price, T. M. and Forbes, D. J. (1987). Inhibition of in vitro nuclear transport by a lectin that binds to nuclear pores. J. Cell Biol. 104, 189–200.

Firmbach-Kraft, I. and Stick, R. (1993). The role of CaaX-dependent modifications in membrane association of Xenopus nuclear lamin B3 during meiosis and the fate of B3 in transfected mitotic cells. J. Cell Biol. 123, 1661–1670.

Foeger, N., Wiesel, N., Lotsch, D., Mücke, N., Kreplak, L., Aebi, U., Gruenbaum, Y. and Herrmann, H. (2006). Solubility properties and specific assembly pathways of the B-type lamin from Caenorhabditis elegans. J. Struct. Biol. 155, 340–350.

Gant, T. M., Harris, C. a and Wilson, K. L. (1999). Roles of LAP2 Proteins in Nuclear Assembly and DNA Replication: Truncated LAP2β Proteins Alter Lamina Assembly, Envelope Formation, Nuclear Size, and DNA Replication Efficiency in Xenopus laevis Extracts. J. Cell Biol. 144, 1083–1096.

Geisterfer, Z. M., Guilloux, G., Gatlin, J. C. and Gibeaux, R. (2021). The Cytoskeleton and Its Roles in Self-Organization Phenomena: Insights from Xenopus Egg Extracts. Cells 10,.

Grange, M., Vasishtan, D. and Grünewald, K. (2017). Cellular electron cryo tomography and in situ sub-volume averaging reveal the context of microtubule-based processes. J. Struct. Biol. 197, 181–190.

Guelen, L., Pagie, L., Brasset, E., Meuleman, W., Faza, M. B., Talhout, W., Eussen, B. H., De Klein, A., Wessels, L., De Laat, W., et al. (2008). Domain organization of human chromosomes revealed by mapping of nuclear lamina interactions. Nature 453, 948–951.

Guo, Y. and Zheng, Y. (2015). Lamins position the nuclear pores and centrosomes by modulating dynein. Mol. Biol. Cell 26, 3379–3389.

Guo, Y., Kim, Y., Shimi, T., Goldman, R. D. and Zheng, Y. (2014). Concentration-dependent lamin assembly and its roles in the localization of other nuclear proteins. Mol. Biol. Cell 25, 1287–1297.

Hampoelz, B., Mackmull, M. T., Machado, P., Ronchi, P., Bui, K. H., Schieber, N., Santarella-Mellwig, R., Necakov, A., Andrés-Pons, A., Philippe, J. M., et al. (2016). Pre-assembled Nuclear Pores Insert into the Nuclear Envelope during Early Development. Cell 166, 664–678.

Heald, R. and Mckeon, F. (1990). Mutation of Phosphorylation Sites in Lamin A That Prevent Nuclear Lamina Disassembly in Mitosis. Cell 61, 579–589.

Heald, R., Tournebize, R., Blank, T., Sandaltzopoulos, R., Becker, P., Hyman, A. and Karsenti, E. (1996). Self-organization of microtubules into bipolar spindles around artificial chromosomes in Xenopus egg extracts. Nature 382, 420–425.

Heitlinger, E., Peter, M., Lustig, A. and Nigg, E. A. (1991). Expression of Chicken Lamin B2 in Escherichia coli: Characterization of its Structure, Assembly, and Molecular Interactions. J. Cell Biol. 113, 485–495.

Herrmann, H. and Aebi, U. (2016). Intermediate filaments: Structure and assembly. Cold Spring Harb. Perspect. Biol. 8,.

Herrmann, H., Häner, M., Brettel, M., Müller, S. A., Goldie, K. N., Fedtke, B., Lustig, A., Franke, W. W. and Aebi, U. (1996). Structure and assembly properties of the intermediate filament protein vimentin: The role of its head, rod and tail domains. J. Mol. Biol. 264, 933–953.

Herrmann, H., Häner, M., Brettel, M., Ku, N. O. and Aebi, U. (1999). Characterization of distinct early assembly units of different intermediate filament proteins. J. Mol. Biol. 286, 1403–1420.

Hetzer, M., Bilbao-corte, D., Walther, T. C., Gruss, O. J. and Mattaj, I. W. (2000). GTP Hydrolysis by Ran Is Required for Nuclear Envelope Assembly. Mol. Cell 5, 1013–1024.

Horn, H. F. (2014). LINC Complex Proteins in Development and Disease. Curr. Top. Dev. Biol. 109, 287–321.

Karabinos, A., Schünemann, J., Meyer, M., Aebi, U. and Weber, K. (2003). The single nuclear lamin of Caenorhabditis elegans forms in vitro stable intermediate filaments and paracrystals with a reduced axial periodicity. J. Mol. Biol. 325, 241–247.

Kittisopikul, M., Shimi, T., Tatli, M., Tran, J. R., Zheng, Y., Medalia, O., Jaqaman, K., Adam, S. A. and Goldman, R. D. (2021). Computational analyses reveal spatial relationships between nuclear pore complexes and specific lamins. J. Cell Biol. 220,.

Lemière, J., Real-Calderon, P., Holt, L. J., Fai, T. G. and Chang, F. (2022). Control of nuclear size by osmotic forces in Schizosaccharomyces pombe. Elife 11,.

Lourim, D. and Krohne, G. (1993). Membrane-Associated Lamins in Xenopus Egg Extracts: Identification of Two Vesicle Populations. J. Cell Biol. 123, 501–512.

Ma, L., Tsai, M. Y., Wang, S., Lu, B., Chen, R., Yates, J. R., Zhu, X. and Zheng, Y. (2009). Requirement for Nudel and dynein for assembly of the lamin B spindle matrix. Nat. Cell Biol. 11, 247–256.

Mall, M., Walter, T., Gorjánácz, M., Davidson, I. F., Ly-Hartig, T. B. N., Ellenberg, J. and Mattaj, I. W. (2012). Mitotic lamin disassembly is triggered by lipid-mediated signaling. J. Cell Biol. 198, 981–990.

Marin, H. C., Allen, C., Simental, E., Martin, E. W., Panning, B., Al-Sady, B. and Buchwalter, A. (2025). The nuclear periphery confers repression on H3K9me2-marked genes and transposons to shape cell fate. Nat. Cell Biol. 27, 1311–1326.

Martins, B., Sorrentino, S., Chung, W. L., Tatli, M., Medalia, O. and Eibauer, M. (2021). Unveiling the polarity of actin filaments by cryo-electron tomography. Structure 29, 488–498.e4.

Meier, E., Miller, B. R. and Forbes, D. J. (1995). Nuclear pore complex assembly studied with a biochemical assay for annulate lamellae formation. J. Cell Biol. 129, 1459–1472.

Mitchison, T. J. (2019). Colloid osmotic parameterization and measurement of subcellular crowding. Mol. Biol. Cell 30, 173–180.

Moir, R. D., Donaldson, A. D. and Stewart, M. (1991). Expression in Escherichia coli of human lamins A and C: influence of head and tail domains on assembly properties and paracrystal formation. J. Cell Sci. 99, 363–372.

Morgan, K. J., Carley, E., Coyne, A. N., Rothstein, J. D., Lusk, C. P. and King, M. C. (2025). Visualizing nuclear pore complex plasticity with pan-expansion microscopy. J. Cell Biol. 224,.

Mücke, N., Kämmerer, L., Winheim, S., Kirmse, R., Krieger, J., Mildenberger, M., Baßler, J., Hurt, E., Goldmann, W. H., Aebi, U., et al. (2018). Assembly Kinetics of Vimentin Tetramers to Unit-Length Filaments: A Stopped-Flow Study. Biophys. J. 114, 2408–2418.

Murray, A. W. (1991). Cell Cycle Extracts. Methods Cell Biol. 36, 581–605.

Nachury, M. V. and Weis, K. (1999). The direction of transport through the nuclear pore can be inverted. Proc. Natl. Acad. Sci. U. S. A. 96, 9622–9627.

Pedersen, R. T. A., Hassinger, J. E., Marchando, P. and Drubin, D. G. (2020). Spatial regulation of clathrin-mediated endocytosis through position-dependent site maturation. J. Cell Biol. 219,.

Rabut, G., Doye, V. and Ellenberg, J. (2004). Mapping the dynamic organization of the nuclear pore complex inside single living cells. Nat. Cell Biol. 6, 1114–1121.

Raghunayakula, S., Subramonian, D., Dasso, M., Kumar, R. and Zhang, X. D. (2015). Molecular characterization and functional analysis of annulate lamellae pore complexes in nuclear transport in mammalian cells. PLoS One 10, 1–27.

Reddy, K. L., Zullo, J. M., Bertolino, E. and Singh, H. (2008). Transcriptional repression mediated by repositioning of genes to the nuclear lamina. Nature 452, 243–247.

Romano, F. B., Blok, N. B. and Rapoport, T. A. (2019). Peroxisome protein import recapitulated in Xenopus egg extracts. J. Cell Biol. 218, 2021–2034.

Sachweh, J., Börmel, M., Klumpe, S., Becker, A., Taniguchi, R., Kubańska, M. A., Pintschovius, V., Kaindl, E., Plitzko, J. M., Wilfling, F., et al. (2025). The small GTPase Ran defines nuclear pore complex asymmetry. Cell 188, 5931–5946.e16.

Sapra, K. T., Qin, Z., Dubrovsky-Gaupp, A., Aebi, U., Müller, D. J., Buehler, M. J. and Medalia, O. (2020). Nonlinear mechanics of lamin filaments and the meshwork topology build an emergent nuclear lamina. Nat. Commun. 11, 1–14.

Segura-Totten, M., Kowalski, A. K., Craigie, R. and Wilson, K. L. (2002). Barrier-to-autointegration factor: Major roles in chromatin decondensation and nuclear assembly. J. Cell Biol. 158, 475–485.

Shi, X., Li, Q., Dai, Z., Tran, A. A., Feng, S., Ramirez, A. D., Lin, Z., Wang, X., Chow, T. T., Chen, J., et al. (2021). Label-retention expansion microscopy. J. Cell Biol. 220,.

Smythe, C., Jenkins, H. E. and Hutchison, C. J. (2000). Incorporation of the nuclear pore basket protein Nup153 into nuclear pore structures is dependent upon lamina assembly: Evidence from cell-free extracts of Xenopus eggs. EMBO Journal 19, 3918–3931.

Tsai, M. Y., Wang, S., Heidinger, J. M., Shumaker, D. K., Adam, S. A., Goldman, R. D. and Zheng, Y. (2006). A Mitotic Lamin B Matrix Induced by RanGTP Required for Spindle Assembly. Science 311, 1887–1893.

Turgay, Y., Eibauer, M., Goldman, A. E., Shimi, T., Khayat, M., Ben-Harush, K., Dubrovsky-Gaupp, A., Sapra, K. T., Goldman, R. D. and Medalia, O. (2017). The molecular architecture of lamins in somatic cells. Nature 543, 261–264.

Tyanova, S., Temu, T., Sinitcyn, P., Carlson, A., Hein, M. Y., Geiger, T., Mann, M. and Cox, J. (2016). The Perseus computational platform for comprehensive analysis of (prote)omics data. Nat. Methods 13, 731–740.

Walther, T. C., Askjaer, P., Gentzel, M., Habermann, A., Griffiths, G., Wilm, M., Mattaj, I. W. and Hetzer, M. (2003). RanGTP mediates nuclear pore complex assembly. Nature 424, 689–694.

Weber, K. and Bement, W. (2002). F-actin serves as a template for cytokeratin organization in cell free extracts. J. Cell Sci. 115, 1373–1382.

Wiese, C., Wilde, A., Moore, M. S., Adam, S. A., Merdes, A. and Zheng, Y. (2001). Role of Importin-β in coupling Ran to downstream targets in microtubule assembly. Science 291, 653–656.

Wiesel, N., Mattout, A., Melcer, S., Melamed-Book, N., Herrmann, H., Medalia, O., Aebi, U. and Gruenbaum, Y. (2007). Laminopathic mutations interfere with the assembly, localization, and dynamics of nuclear lamins. Proc. Natl. Acad. Sci. U. S. A. 105, 180–185.

Wilde, A. and Zheng, Y. (1999). Stimulation of microtubule aster formation and spindle assembly by the small GTPase Ran. Science 284, 1359–1362.

Zhang, C. and Clarke, P. R. (2000). Chromatin-Independent Nuclear Envelope Assembly Induced by Ran GTPase in Xenopus Egg Extracts. Science 288, 1429–1432.

Zhang, C., Jenkins, H., Goldberg, M. W., Allen, T. D. and Hutchison, C. J. (1996). Nuclear lamina and nuclear matrix organization in sperm pronuclei assembled in Xenopus egg extract. J. Cell Sci. 109, 2275–2286.

Zheng, X., Hu, J., Yue, S., Kristiani, L., Kim, M., Sauria, M., Taylor, J., Kim, Y. and Zheng, Y. (2018). Lamins Organize the Global Three-Dimensional Genome from the Nuclear Periphery. Mol. Cell 71, 802–815.e7.

Zhuang, Y. and Shi, X. (2023). Expansion microscopy: A chemical approach for super-resolution microscopy. Curr. Opin. Struct. Biol. 81, 102614.

Zhuang, Y. and Shi, X. (2024). Label-Retention Expansion Microscopy (LR-ExM) for Enhanced Fluorescent Signals using Trifunctional Probes. Curr. Protoc. 4, 1–13.

